# Centripetal nuclear shape fluctuations associate with chromatin condensation towards mitosis

**DOI:** 10.1101/2021.11.25.469847

**Authors:** Viola Introini, Gururaj Rao Kidiyoor, Giancarlo Porcella, Marco Foiani, Pietro Cicuta, Marco Cosentino Lagomarsino

## Abstract

The cell nucleus plays a central role in several key cellular processes, including chromosome organisation, replication and transcription. Recent work intriguingly suggests an association between nuclear mechanics and cell-cycle progression, but many aspects of this connection remain unexplored. Here, by monitoring nuclear shape fluctuations at different cell cycle stages, we uncover increasing inward fluctuations in late G2 and early mitosis, which are initially transient, but develop into instabilities that culminate into nuclear-envelope breakdown in mitosis. Perturbation experiments and correlation analysis reveal an association of these processes with chromatin condensation. We propose that the contrasting forces between an extensile stress and centripetal pulling from chromatin condensation could link mechanically chromosome condensation and nuclear-envelope breakdown, the two main nuclear processes during mitosis.

**Significance Statement:** The nucleus was recently shown to exhibit shape fluctuations that vary with cell-cycle stage, but we know very little about the possible links between nuclear mechanics and cell cycle- progression. Through flickering analysis, this study monitors radius and nuclear envelope fluctuations across the cell cycle. The authors discover that as the cell cycle progresses towards mitosis, localised inward invaginations of the nuclear shape form initially transiently and gradually increasing their amplitude, in association with chromatin condensation. This phenomenon develops into nuclear envelope breakdown, suggesting a novel link between cell cycle, chromatin mechanics and nuclear shape fluctuations.

---

**T**he shape fluctuations (also known as flickering) of vesicles *in vitro*, are driven by thermal motion, to which the membranes respond passively. Specifically, the observed transient shape fluctuations can be interpreted in terms of equilibrium states (1), and their measure by time-lapse microscopy provides a powerful biophysical tool to characterise the constituent mechanical properties: parameters such as bending modulus, tension and viscosities (2). The validity of these tools, and the assumption of thermal equilibrium, can extend to some living systems where the action of molecular motors and other active forces are not relevant, such as erythrocytes (3, 4).

In more complex living systems, chemical energy is turned into mechanical forces, e.g. by molecular motors, and these forces add to the thermal forces to induce fluctuations of cells and cellular compartments (5). In this scenario, the extent of shape change is related to both the level of non-thermal (nonequilibrium) activity and to the ‘active’ mechanics. Teasing out these two contributions is very difficult and has been achieved only in a few cases (6, 7). It requires performing deformation assays under the presence of a known external force, and comparing the outcome to the spontaneous deformations. In a living cell or tissue, the active (ATP-driven) forces enhance shape fluctuations by non-equilibrium pulling and stress-relaxation events (e.g. through pumps and cytoskeletal elements such as actomyosin).

Abundant evidence shows that nonequilibrium processes can drive membrane flickering to more complex behavior than predicted by thermodynamics equilibrium, for example causing a breakdown of the “fluctuation-dissipation” theorem valid at equilibrium (7), which links the decay of spontaneous fluctuations to the response to external perturbations. In such conditions, monitoring shape fluctuations is still useful, but the precise identification of biophysical parameters such as tension or stiffness is more difficult, and one can generally refer to “effective” (or “apparent”) tension and bending moduli as a complex byproduct of passive membrane properties and the result of active driving forces. In such conditions the equilibrium model may still be a useful guide, for example allowing to compare the relative amplitudes of fluctuations at different wavelengths. In many cases, the active fluctuations can be reduced to the standard model with an “effective temperature”, by which the active forces simply increase the noise level with respect to the thermal motion (8).

The cell nucleus shows a complex shape and size dynamics during the cell cycle (9–12). It is confined within the nuclear envelope (NE), a complex quasi-two-dimensional structure comprising two lipid bilayer membranes separated by a perinuclear space of 20-40 nm and mechanically linked nuclear lamina, a 50-80 nm thick network formed by lamins (11). The cytoskeleton and chromatin maintain direct links with the NE and thus between themselves through the LINC complex and other linker proteins. This mechanical coupling of cytoskeleton and chromatin enables the transmission of external mechanical cues across the NE (11) via lamina-associated domains to the chromatin, thereby regulating transcriptional activity and other nuclear processes (13, 14).

Recently, Chu and coworkers have monitored nuclear undulations in mammalian cells at timescales of seconds, revealing cell-cycle dependent flickering (15). These undulations are likely actively driven both internally by the nucleus (as evidenced by an increase of undulations upon inhibition of transcription) and externally by the cytoskeleton (evidenced by biochemical perturbations of actomyosin). The authors have hypothesized that regulating flickering may aid and tune nuclear transport through nuclear pores, yet many questions on the connection of cell-cycle dependent nuclear shape fluctuations and cell-cycle progression remain open, in particular regarding a possible role played by nuclear mechanics in the cell cycle itself. Indeed, emerging evidence indicates that in cycling cells both external and internal mechanical forces trigger important changes in nuclear structure, activity and composition (10, 11, 16). Additionally, recent studies show a link between nuclear mechanics and cell cycle progression in cancer cells and epithelia, in particular linking nuclear tension to the G1/S transition (17, 18). Finally, the nucleus was reported to act as a ‘ruler’ in cells moving through constrictions, which rely on nuclear mechano-signalling to modulate forces enabling their passage through restrictive pores (by mechanically coupled signaling of the cPLA2 protein) (19). Intriguingly, cells in G2 appear to have a larger such ruler, hence require less confinement than G1 cells to trigger the contractile response. Consequently, the hypothesis was formulated that these cues could couple the cell mechanical environment to cell-cycle progression.

Here, we analyse NE fluctuations by high-resolution video microscopy, testing the connection between nuclear mechanics and cell-cycle progression, particularly focusing on the transition from G2 phase to the onset of mitosis, finding increasing transient inwards deformations that associate to chromatin condensation.

## Results

### Nuclear shape fluctuations vary with the cell cycle

We first tested whether our cells showed a similar cell-cycle dependency of the NE flickering as observed by Chu *et al*. We used a GFP- tagged version of the Emerin protein to mark the NE. Since it is well known that Lamin over-expression can directly impact mechanical properties of the nucleus (20), we decided to use a tagged version of Emerin instead of Lamin to minimise the adverse effects of labeling on nuclear behavior. Analysing the cell cycle duration of the HeLa cells by performing time-lapse videos (see SI) showed that on average, in our growth conditions, HeLa cells spent 16.48 hours in interphase and 1.75 hours undergoing mitosis (**SI Fig. S1a**), consistent with previous reports (21). Nuclear area increased throughout interphase, then decreased with entry to mitosis (**SI Fig. S1b**). Cells were arrested at the G1/S transition (by a double thymidine block) or at the G2/M transition (by CDK1 inhibition) and then released to follow them through cell cycle progression. Cell cycle stage was univocally assigned by monitoring the cells every 3 hours from release (**Fig. 1a**).

**Fig. 1.**
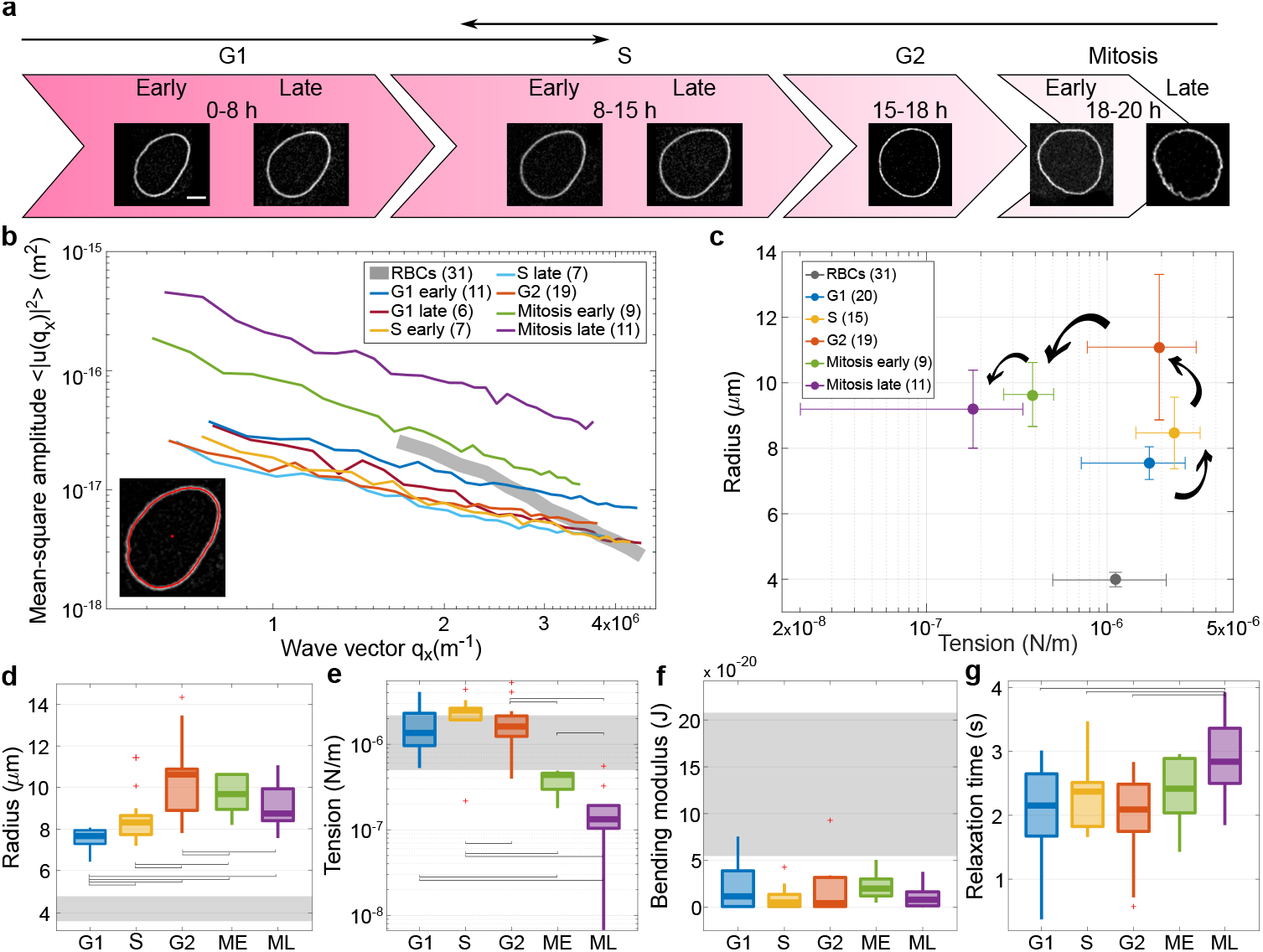
Shape fluctuations of HeLa cell nuclei are cell-cycle dependent and increase during mitosis. (a) Snapshots of a representative nucleus at 7 time points from the start at early G1 phase, throughout S, G2 and mitosis (scale bar 5 *µ*m). Arrowheads indicate the reference time-points to determine cell cycle phase. (b) Average spectra of wave vector-dependent fluctuation amplitudes (modes 6-34) for cells at different stages in the cell cycle. The number of nuclei considered for each cell-cycle stage are reported in the legend in brackets. The fluctuation amplitude 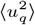 exhibits a decrease with increasing time from G1 until G2, where the fluctuations are reduced by about 3 times. Instead, active nuclear fluctuations during mitosis become 4 times higher in early mitosis (green line) and 10 times in late mitosis (purple line). Inset: contour detection of NE (red line) with fluorescent label Emerin. The initial manual selection of the center (red dot) and an initial point on the NE define the annular region containing the cell boundary used in image analysis. (c) Effective tension *vs* radius scatterplot shows clusters from different cell-cycle stages forming an open counterclockwise trajectory. (d-f) Box plots of shape-fluctuation parameters throughout the cell cycle (significance indicated. The data show no significant changes (p value > 0.05) in effective bending modulus across the cell cycle, while effective tension increases significantly during S phase and decreases up to one order of magnitude during mitosis. The cell radius increases from the starting point in G1 until G2 and then does not change much. (g) the characteristic relaxation time for mode 3 becomes longer during mitosis. Grey bands and markers represent RBC fluctuation parameters. P values are reported in **SI Table S2**; significant relations are highlighted with brackets.

**Fig. 1b** shows the average fluctuation spectra at different stages of cell cycle, i.e., the amplitudes of fluctuations, calculated by the deviation of the instantaneous contour from the average contour for all the recording frames, and plotted as a function of wave vector (inverse wavelength of the projected shape deformation) (22). Nuclear fluctuations decrease by about three fold from early G1 to late S phase (**SI Video 1**), which is in line with data shown by Chu and coworkers after 13 hours of release from mitosis arrest (15). However, we also report a notable increase of these fluctuations already in late G2 and early phases of mitosis, which develops into dramatic deformations during mitosis when the cells start to round up, eventually developing into an instability triggering NE breakdown (**SI Video 2, Fig. 1b**).

Although the system is out of equilibrium, if we assume that the active forces play the role of an increased ‘effective temperature’ then it is possible to use the standard model for fluctuations (15), and extract effective biophysical parameters. As anticipated above, it is important to stress that these measured effective parameters are not the same as the biophysical ones but a byproduct of constitutive parameters and the action of active forces. We will refer to these as effective tension and bending modulus in the following, and explicitly discuss their interpretation whenever necessary.

Compared to previous work, we adopted two important technical improvements. First, we account for the projection of fluctuations on the equator in the measured shape deformations (3, 22, 23), which were neglected in Chu *et al*. work and lead to erroneous *q* dependencies. Second, we consider spectra as a function of wave vector rather than wave number, in order not to average together fluctuations of different wavelength from nuclei of different size. We obtain the average square displacement 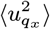, where the wave vector of the projected equatorial profile is *q*_*x*_ = 2*πn/L, L* is the length of the profile and *n* = 0, 1, 2, … the modes (3). Effective bending modulus and tension are then obtained by a fit of the spectra with the formula

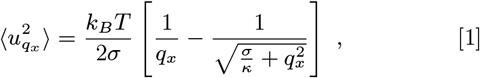

where *σ* is an effective tension (measuring, in the passive case, mechanical response to extensile stress) and *κ* is a an effective bending rigidity (measuring response to curvature in the passive case). *k*_*B*_ is Boltzmann’s constant, and *T* is the absolute temperature. By taking *T* as the physical temperature, we interpret any non-equilibrium effect within the parameters *σ* and *κ*. This seems justified here because the mode-dependence of the data is consistent with the equilibrium model. Considering an active surface like the NE with this model, *σ* can be interpreted as the resistance of the surface to change total area, in response to active and thermal forces, and *κ* as the total energy necessary to bend and ruffle the surface. (8, 24).

Equation (1) has limiting behaviors 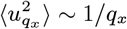 for modes dominated by effective membrane tension, and 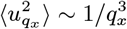 for modes dominated by effective bending rigidity. For the range of modes considered in our analysis, the nuclear fluctuations are mainly affected by effective tension, as the mean-square amplitude spectrum is dominated by the 1*/q*_*x*_ trend (4, 22).

It is instructive to plot nuclear radius versus effective tension in the different cell-cycle stages (**Fig. 1c**). Effective tension initially increases with radius, as would be expected for an “inflated” passive membrane. However, this trend is inverted starting from the G2 phase, so that radius keeps increasing while effective tension is reduced. The physical properties of the lamina may change significantly in this part of the cell cycle due to lamin phosphorilation (25, 26), but an increase in the active forces could also concomitantly drive nuclear shape fluctuations. Hence the decrease in effective tension in late G2 and mitosis could be due to a drop in physical tension, and/or an effect of active forces. Overall, **Fig. 1c** shows how during the cell cycle the nucleus follows a counterclockwise trajectory in the effective tension-radius plane, which starts at nucleus birth and culminates in NE breakdown at late mitosis. The changes in radius and effective tension across phases of cell cycle are statistically significant (**Fig. 1d-f**), while effective bending rigidity, remains fairly constant. Looking at cells that were imaged over several stages of the cell cycle, we verified which of the average trends of the parameters were robust in single cell trajectories (**SI Fig. S2**).

From our measurements, it is possible to monitor the relaxation time scales of the dominant deformation modes. For a passive membrane, the modes decay exponentially and the relaxation time scale is the ration of the modulus driving the relaxation, and the viscosity. When the modulus is determined from a passive spectrum, e.g. the tension, then this allows to determine the viscosity. However, for an active surface such as this one, the time scales reflect active dynamics, and the decay of the modes can become very complex. We considered the relaxation time *τ* of mode 3, where we found that no complex behavior appears and the decay is a simple exponential, (see **SI Fig. S3**). **Fig. 1g** displays longer relaxation time for late mitosis. This can be interpreted as a signature of active fluctuations/deformations from active nuclear or cytoplasmic pulling or pushing elements, visible in the movies during mitosis, which could trigger different characteristic times (due to the dynamics of the active elements) than those of passive relaxation.

Red blood cell (RBC) fluctuations have been extensively studied, representing a simpler well-understood system, yet with some common biophysical properties in common with cell nuclei (e.g., being supported by cytoskeletal elements). Hence, we decided that it could be instructive to use them as a reference, and we compared the behavior of HeLa cell nuclei with those of RBC (grey bands in **Fig. 1**). HeLa nuclei have in general larger dimensions, a longer relaxation time, and smaller effective bending modulus, but their effective tension is similar to RBC if we exclude the dramatic changes occurring for nuclei at mitosis (3, 4). The mean and SEM of nuclear biophysical properties from HeLa cells at different stages of the cell cycle are reported in **SI Table S1**, and p values in **Table S2**.

### Calyculin A treatment recapitulates the behavior of nuclear shape fluctuations during mitosis

Mitosis is a complex mechanochemical process requiring coordinated action from multiple cellular components mediated by several kinases and signalling molecules. The cellular shape alterations at the beginning of mitosis are accompanied by chromosome condensation and dramatic remodelling of the cortical actin cytoskeleton (12), which eventually lead to NE breakdown. We wondered to what extent all these processes could be directly linked to nuclear-shape behavior and, particularly what is the role of chromatin condensation. This hypothesis would be supported if shape-deformation behavior of late-G2 and mitotic nuclei could be reproduced either by inducing chromatin condensation or perturbing the actin cytoskeleton. Therefore, in order to correlate chromatin dynamics with nuclear shape deformations, and to disconnect the shape fluctuations in mitotic cells to mitosis-specific chemical signalling, we treated cells with calyculin A, a drug that induces rapid premature chromatin condensation in all cells independent of their cell cycle (27, 28). Shape fluctuations of calyculin-A treated nuclei were recorded and compared with the fluctuations of the same cell observed prior to drug treatment (**SI Video 3**). Next, we wanted to understand better the individual contributions of condensing chromatin and de-polymerising actin in mitotic nuclei shape fluctuation phenomenon. Therefore, we treated G2/M arrested cells with actin de-polymerising drug latrunculin A, but in presence of G2/M arresting drug (R3306). This allowed us to disrupt actin cytoskeleton without inducing chromatin condensation in late G2 cells, which otherwise would have entered into mitosis (**SI Video 4**).

**Fig. 2a** shows that after short exposure to calyculin A (20 min) and latrunculin A (25 min), the effective tension of nuclei is reduced in a similar fashion to what happens in the G2-mitosis transition (G2 phase is the control). A longer (50 min) exposure of cells to latrunculin A leaves nuclear radius and effective tension constant. Calyculin A instead produces within the same treatment time, a subpopulation of cells with a further reduction of nuclear radius and a further decrease in effective tension, resembling late mitosis. Specifically, **Fig. 2b and c** compare the radii and effective tension of interphase and mitotic nuclei, with the same nuclei before and after treatment with calyculin A and latrunculin A.

**Fig. 2.**
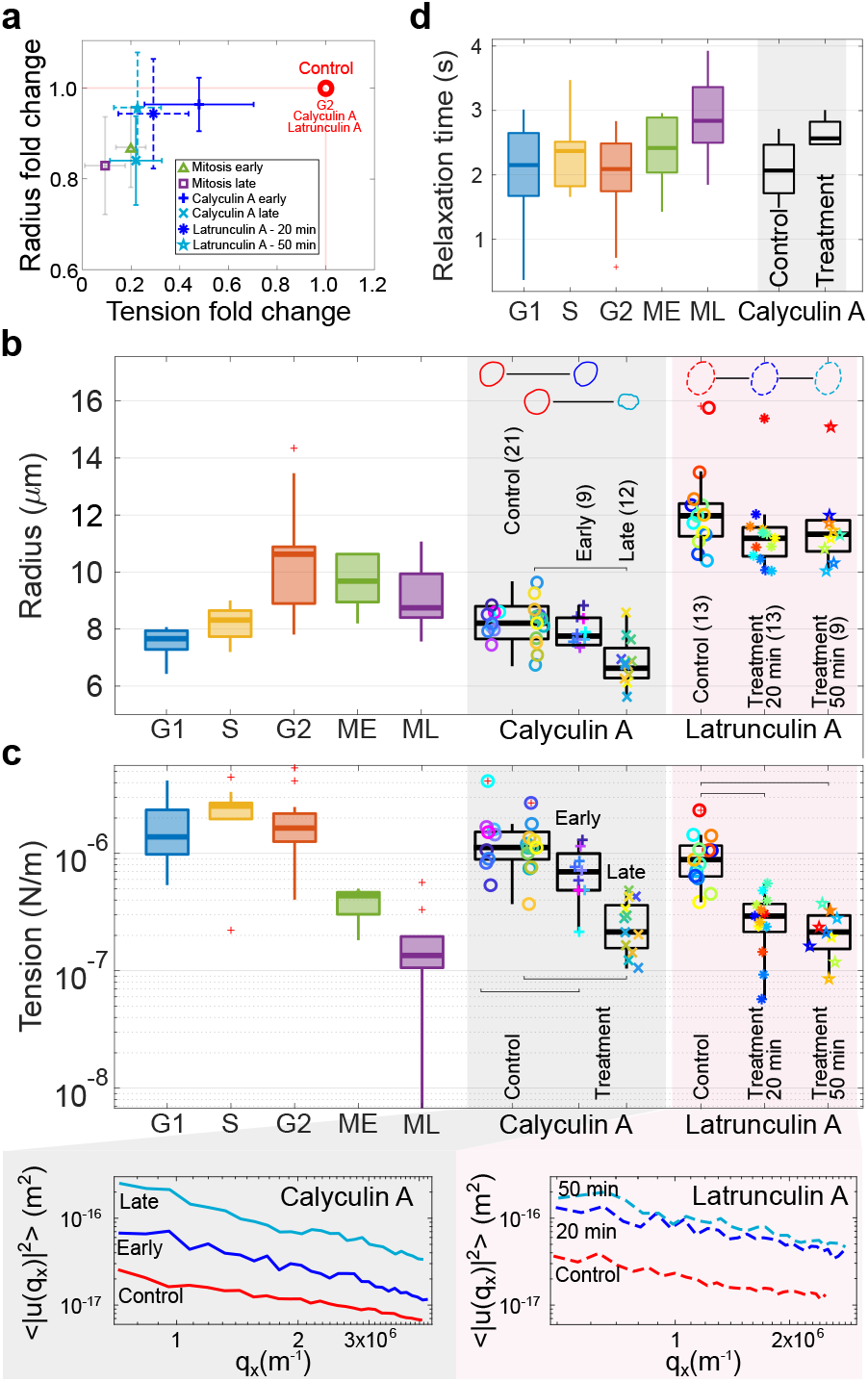
Calyculin A and latrunculin A treatments recapitulate the joint radius / effective tension changes found in mitosis. (a) Radius-tension change after treatments compared to the control phase G2, and early and late mitosis for cycling cells. Calyculin A causes a reduction of both radius and effective tension, while latrunculin A decreases effective tension to a lower value, which remains constant with treatment time (early, 25 min *vs* late, 50 min). (b-c) Details of radius and tension of the same cells before and after both treatments, compared to the values throughout the cell cycle. Shape changes of representative nuclei are highlighted in panel b. The insets below panel (c) report the respective averaged fluctuation spectra. (d) Relaxation time of mode 3 slightly increases after calyculin treatment, as for mitosis. P values are reported in **SI Table S2**; significant relations are highlighted with brackets.

In fact, approximately 10-15 minutes after calyculin A treatment, nuclei start showing shape fluctuations similar to early-stage mitotic nuclei (henceforth termed “early calyculin A”) without a major deformation of the nuclear shape. These fluctuations, within another 2-3 minutes, evolve into widespread invaginations as the cells start rounding up (see **SI Video 3**). These rounded cells resemble the late stage of mitosis, henceforth termed “late calyculin A”, showing significantly distorted nuclear shape compared to their untreated state, as well as reduced nuclear radius. In practice, since the full time lapse was not available for all cells, “early” and “late” calyculin A cells were defined based on nuclear radius changes compared to pre-treatment pictures (see methods). Nuclei included in the latter category show a reduction of radius after treatment of at least 10%. Sketches of nuclei in **Fig. 2b** report typical changes in nuclear shapes after treatment, and further characterisation will be described in **Fig. 3a**. Finally, **Fig. 2d** shows that the relaxation time of mode 3 increases upon calyculin treatment, similarly to mitosis, while it is not affected by latrunculin A (data not shown). We found that calyculin A treatment recapitulates the behavior of mitotic nuclei close to NE breakdown. From this set of data, we conclude that the observed shape fluctuations of mitotic nuclei is mechanically driven by condensing chromatin and cytoskeletal remodelling, therefore mechanical or biochemical signalling triggered at chromatin condensation might be sufficient to generate the nuclear shape remodeling observed before NE breakdown. To address the possible concern that calyculin may affect lamin phosphorilation - which is a signaling part of nuclear envelope breakdown, we performed a Western blot analysis of phospho-Lamin staining in calyculin A treated cells (**SI Fig. S4**), finding no visible change.

**Fig. 3.**
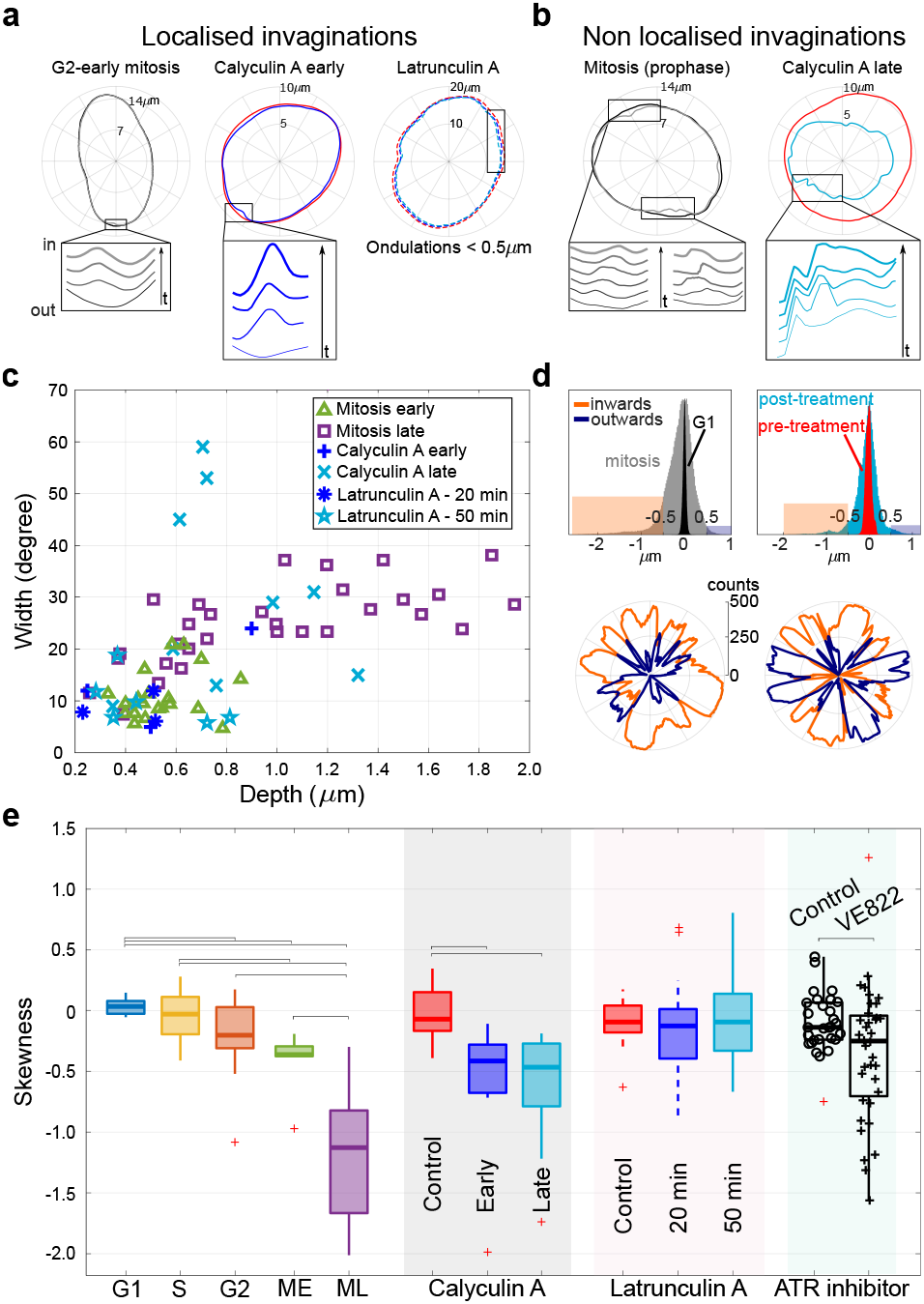
Late-G2 and mitotic deformations are dominated by inwards invaginations, compatible with the action of centripetal pinning forces. (a) Representative examples of the localised inward invaginations that emerge in early mitosis phase and in early stages of calyculin A treatment, but are not found with latrunculin A treatment. (b) Examples of the non-localised invaginations observed in late mitosis and later stages of calyculin A treatment. The insets in panel a and b illustrate the dynamics by snapshots at equal time lags (**SI Videos 2-4**) (c) Invagination width at the maximal deformation increases by 2-3 fold in late mitosis while depth can increase up to 10 fold. Invaginations from early and late phases of calyculin A treatment resemble the ones in mitosis, while latrunculin A treatment has mild effects on the invaginations (they remain within < 1*µ*m in depth and 25 degrees in width). (d) The histograms (top) as well as polar plots (bottom) of signed shape fluctuations show the bias towards inward motion of mitosis and calyculin A late nuclei (orange are inward and blue outward fluctuations). Histograms count all contour angles for 500 frames; inward fluctuations were considered < -0.5*µ*m (orange band),and outward > 0.5*µ*m (blue band). (e) Boxplots of the skewness of the signed shape fluctuation histograms (shown in panel d) over cell-cycle stages and upon drug treatment. The centripetal asymmetry increases during mitosis and calyculin A treatment, while latrunculin A does not affect it. ATR inhibitor VE822 increases the events with negative skewness. P-values are reported in **SI Table S2**; significant relations are highlighted with brackets.

Since calyculin A activates myosin-2 mediated contractility (29), we checked whether the increased centripetal invaginations observed upon treatment with this drug could be a byproduct of increased actomyosin contractility. As a control, we performed a double chemical perturbation with calyculin A and blebbistatin (a myosin inhibitor). Nuclear fluctuations, as well as their radius and effective tension, replicate the nuclear features after treatment with calyculin A. Treatment with blebbistatin alone did not affect the dominance of inwards *vs* outwards fluctuations, and, coherently with previous reports (15), increased effective tension (**SI Video 5, SI Fig. S5**). Since blebbistatin is known to be inactivated in blue light, we confirmed the result with Y27632 (ROCK inhibitor), a rho kinase inhibitor that decreases actomyosin contractility and is not affected by illumination (**SI Fig. S5**). Likely the effects of the low illumination used in our experiments on blebbistatin are mild or absent, and we report identical phenotype as in. (15). Taken together, these results support the interpretation that calyculin-induced nuclear shape deformations are not due to increased actomyosin contraction.

Finally, we found that actin depolymerization increases shape fluctuations, but does not affect nuclear size. Unlike calyculin A treated nuclei, nuclei of latrunculin A-treated cells develop shape fluctuations within 20 min of treatment, resembling the behavior of early mitotic nuclei. However these fluctuations do not progress with time into more prominent and irreversible invaginations and shape alterations, even after 50 min of drug treatment (**Fig. 2c**). The mean and SEM of nuclear biophysical properties from cells upon calyculin A and latrunculin A perturbations are reported in **SI Table S3** together with their statistically different p values (**SI Table S2**).

### Mitotic fluctuations are invaginations mediated by centripetal pinning forces

We noticed that most of the transient deformations contributing to the decrease in effective tension from G2 to mitosis had two specific properties: (i) they were localised in one region of the observed profiles and (ii) they looked like the tip of the deformation pointed towards the inner side of the nucleus (**SI Video 2**). During late mitosis, the deformations became more widespread, and increased dramatically their asymmetry towards the nucleus center. As the cells progressed towards NE breakdown, we observed that the inward deformations became increasingly long-lived and less localised, as increasingly larger “patches” of the lamina appeared to be displaced centripetally. Eventually, these deformations became unstable, and instead of being restored to an equilibrium shape, they developed into the deformations leading to NE breakdown. **Fig. 3a and b** confirm this behavior, which was also found in early and late stage calyculin-A treated cells. Conversely, latrunculin A treatment does not cause invaginations, although reducing nuclear effective tension. During late mitosis and late stage calyculin A-treated nuclei, invaginations become wider and deeper (**Fig. 3c**). Invagination width at the maximal deformation increases by 2-3 fold in late mitosis, and depth up to 10 fold. Nuclear invaginations from early and late phases of calyculin A treatment follow the trend of invaginations progressing through mitotis in untreated cells, as opposed to latrunculin A treatment, where invaginations remain within less than 1*µ*m in depth and 25 degrees in width. Depth was calculated as the difference between the steady state contour and the minimum of the invagination, and the width by the points corresponding to 10% of the depth. To characterise such inward deformations, we considered the distribution of *signed* shape fluctuations, defined as the integrated difference between the profile and a reference profile calculated as the average shape of ten frames before the invagination developed (**Fig. 3d,e**). Inward invaginations (< -0.5*µ*, orange band in **Fig. 3d**) are prevalent with respect to outwards (> 0.5*µ*m, blue band), shown both in the histograms and relative polar plots. Distribution of fluctuations for all frames and angles are wider for cell mitosis (grey histogram in **Fig. 3d**) and post-calyculin A treatment (cyan), with respect to interphase (G1 and pre-treatment respectively black and red histograms). Polar plots represent the total number of frames of inward and outward fluctuations at every angle.

To quantify the behavior of inwards vs outwards deformations, **Fig. 2e** uses the skewness of their distribution as summary statistics. A negative skewness corresponds to an enrichment in inward deformations. The results confirm that inward deformations increase in early and late mitosis, as well as upon calyculin A treatment, while it is unaffected by latrunculin A. Incidentally, we noticed that the typical shape of inward deformations fits well the theoretical shape of a membrane deformed by a localised pinning force (30), as reported in **SI Fig. S6**. However, our lack of knowledge of the actual biophysical parameters (bending rigidity and tension) prevents us from using this shape to estimate the magnitude of the pulling forces.

### Chromatin density increases in correspondence to centripetal nuclear shape deformations

As a last step to gain further insight into a possible role of chromatin in causing the observed centripetal shape fluctuations, we analysed movies of cells in which H2B histones were labelled (with fluorescent m-Cherry) jointly with Emerin (as above). These experiments enabled us to monitor NE shape jointly with chromatin density. We first evaluated whether nuclei in early and late mitosis could exhibit any separation between chromatin and lamina (as proxied by Emerin) at the site of invagination, and we observed no separation for all the cells analyzed (an illustrative example is shown in (**SI Fig. S7**).

Subsequently, we quantified the cross-correlation between local deformations of NE and fluorescence intensity from histones in the corresponding area during invaginations (examples in **SI Fig. S8, SI Video 8**). We observed that in the neighborhood of invaginations inside the nucleus, while NE contour decreases at the the angle of maximum inward pulling, the mean fluorescence intensity of chromatin increases. This observation clearly establishes a link between chromatin state and nuclear shape deformations in case of local reversible invaginations. In some cases (e.g. the second case reported in **SI Fig. S8**), we saw that the chromatin signal increases a few seconds before the inward NE deformation reaches its maximum extent, suggesting that the local chromatin condensation leads to an increase of fluorescence that occurs before the NE invagination, and possibly pulling the NE itself inward.

### ATR inhibitors increase asymmetric deformations and effective bending modulus, and decrease effective tension

The ATR signalling protein, a kinase mostly known as an activator of the DNA damage checkpoint, was reported to be a sensor of mechanical stress, to localise at the nuclear envelope and link it to chromatin, and to facilitate release of chromatin from the NE (31). Since ATR inhibition/depletion causes accumulation of NE invaginations attached to semi-condensed chromatin (31, 32), we reasoned that ATR might trigger the relaxation of the invaginations caused by chromatin condensation by releasing chromatin from the NE. Under this hypothesis, symmetry and extent of nuclear flickering in late G2 to mitosis could be affected by ATR inhibition. In line with these assumptions, **Fig. 3e** shows that ATR inhibition causes an increase in the number of deformation showing negative skewness. This change is not observed for all cells, and this could be interpreted as a cell-cycle-dependent effect of ATR inhibition. Indeed, the ATR effect on membrane fluctuations is expected to be variable with the exact cell-cycle stage (not completely controlled in our experiments) and the time window of inhibition. Despite of these limitations in our experiments, our observations are consistent with more long-lasting centripetal invaginations when ATR is inhibited. The same experiments show a mild decrease in effective tension (in line with our hypothesis) and an increase in effective bending modulus (**SI Fig. S9, SI Video 9**). The measured bending modulus increase in ATR inhibited nuclei is a consequence of the drug having opposite effects on the small-*q* (enhanced) and high-*q* (depressed) part of the fluctuation spectrum, and might be the result of a previously reported change in the lipid composition of the NE upon ATR inhibition (32).

## Discussion

**SI Fig. S10 and SI Video 9** report kymographs that summarize all our main results visually. Our results confirm the scenario proposed by Chu and coworkers (15), whereby nuclear shape fluctuations are driven by a combination of thermal motion and forces from chromatin and cytoskeleton. Fully in line with this study, we find that latrunculin A (actin depolymerization) increases shape fluctuations (decreasing effective tension), and decreases nuclear radius, while blebbistatin (Myosin-II inhibition) increases nuclear radius and effective tension. This data suggests that the dynamic flickering of nuclear envelope might be countered by the presence of actin stress fibers (which are lost with latrunculin A), possibly via LINC connections, while the dynamic rearrangement of stress fibers caused by loss of myosin contractility has a more complex “stiffening” effect, which also (surprisingly) leads to radius increase. In addition to this, Chu *et al*. reported that nuclear processes, including transcription and nuclear transport, also influence nuclear shape fluctuations (**Table 1**). Combining the two observations, we confirm the picture of a NE that is a dynamic component rather than a static organelle, which responds to cellular and nuclear events. However, when considering cell-cycle changes, Chu *et al*. only reported a decrease in amplitude of symmetric fluctuations, with the progress of interphase (G1-S-G2). They interpreted this as a change of material properties and/or a reduction of the forces driving the shape fluctuations. While our observations are compatible with this study, we focused on deformations occurring during G2 phase and onset of mitosis, using different perturbations, which lead us to surprising results.

**Table 1.** Comparison of nuclear-shape fluctuation behavior before mitosis and under biochemical perturbations.

| Perturbations | Biological effect | Fluctuations | Implications in shape fluctuations | Nucleus morphology | References |
| --- | --- | --- | --- | --- | --- |
| Mitosis | Chromatin condensation | Increase (T&A) | Chromatin and cytoskeletal activity | More spherical | This study |
| Calyculin A | Induce chromatin condensation | Increase (A) | Chromatin involvement | Spherical and softer | This study |
| Latrunculin A | Inhibition of actin polymerization | Increase (A) | Cytoskeletal activity | Softer | This study, (15) |
| α-amanitin | Inhibition of polymerase II transcriptional activity | Increase (T&A) | Chromatin involvement | | (15) |
| ATP depletion | Influence transcription, DNA replication, DNA repair, chromatin remodeling | Decrease (T&A) | Active (> 4 s) and passive (> 1s) fluctuations |  | (15) |
| Blebbistatin / Y27632 | Inhibition of myosin II activity | Decrease (T&A) | Myosin II activity contribution | Slightly bigger | This study, (15) |
| Nocodazole | Inhibition of microtubule polymerization | Decrease (T&A) | Microtubules contribution |  | (15) |
| ATR inhibitor | Reduced release of chromatin-envelope link | Increase (T&A) | Chromatin involvement | Invaginations micronuclei | This study, (32) |

Legend: T = thermally driven and A = active. Data from Fig. 1 and 2, and ref. (15).

Specifically, after the genome has completed replication, we find evidence supporting active pinning centripetal forces that drive increasingly strong shape fluctuations (also resulting in a drop in effective tension) from G2 to mitosis, up until NE breakdown. Hence, (i) shape fluctuations can dramatically increase from G2 to mitosis, and (ii) they can become highly non-symmetric at this stage. Fluctuation asymmetry favoring in-wards displacements appears already in G2, together with reversible “pinning” centripetal deformations. These deformations become increasingly long lasting and irreversible as the cell cycle progresses towards NE breakdown. Interestingly, neither latrunculin A nor blebbistatin / Y27632 treatments affect the asymmetry of the shape fluctuations, while calyculin A treatment makes them centripetal. These observations suggest that chromatin dynamics can be related to NE centripetal shape fluctuations.

Our data also allow us to formulate some hypotheses on the force balance between the physical processes that regulate nuclear mechanics. Physically, nuclear shape is set by three mechanical components: chromatin, lamins, and the cytoskeleton. Chromatin and lamin A are typically seen as resistive elements that together maintain nuclear shape. Lamins alone, on the contrary, cannot maintain nuclear shape, and the lamina buckles under mechanical stress when it is unsupported by chromatin, suggesting a physical model of the nucleus as a polymeric shell enclosing a stiffer chromatin gel (14). The role of the cytoskeleton is less clear, and sometimes it is pictured as a compressive force, but a perinuclear actin cap has also been shown to stabilize nuclear shape (33). Disconnecting chromatin from the inner nuclear membrane results in softer nuclei that are deformable and more responsive to cytoskeletal forces (13)

On the basis of our experiments, we formulate the hypothesis of a nucleus under extensile and/or stabilizing stress from the external cytoskeleton, so that condensing chromatin can locally exert inward pulling forces. The calyculin treatment, and the direct joint imaging of H2B histones and Emerin lead us to surmise that these local pinning forces (becoming more widespread as G2 progresses to mitosis) may come from condensing chromatin. This hypothesis deviates from the standard view whereby a chromatin gel confers structural integrity and stiffness to the nucleus, but it does so only in the idea that this gel would be under a pre-stressed condition (14). The decrease in effective tension under latrunculin A treatment is compatible with extensile stress applied by the cytoskeleton. Therefore, our data may support a scenario of fairly uniform extensile stress applied by the cytoskeleton, counterbalanced by local centripetal pulling applied by chromatin.

Condensing chromatin exists in mechanically stressed state (34). The idea that chromatin condensation could alter NE shape by exerting centripetal forces was suggested by previous observations on Drosophila salivary glands (35), where chromatin compaction forces were shown to drive distortions of the NE through chromatin-envelope interactions. Kumar and coworkers showed that chromatin-envelope interactions generate mechanical stress, which recruits and activates ATR kinase at the NE (31). In line with their results, we observe a more negative skewness in nuclear shape deformations when ATR is inhibited near prophase. Additionally, a study on non-tumorigenic mammary epithelial cell MCF-10A has implicated chromatin in nuclear shape deformations, showing that these were independent of cytoskeletal connections (36). Chromatin decompaction was also shown to cause nuclear blebbing, regardless of lamin, as well as nuclear swelling (37, 38). In addition to these findings, our data suggest that the force balance at the NE is not static, and the nucleus progressing from S phase to G2 and mitosis feels constant or increasing extensile stress from the outer cytoskeleton, and increasing localised stress from inner chromatin, affecting its shape fluctuations. Isotropic contributions to these stresses also likely come from forces of osmotic origin (39, 40).

The phenomena observed here could play a role in the coordination of chromatin condensation and NE breakdown during mitosis. It seems natural to think that the timing of these two events should be coordinated - just like NE reassembly should be coordinated with chromosome segregation (10). Since the NE is squeezed between the cell cytoskeleton on the cytoplasmic side and chromatin on the nucleoplasmic side, and both of these active systems undergo major rearrangements over the cell cycle, it is possible that the cell-cycle dependent flickering may be not only a byproduct but also a driver of cell-cycle progression. Since chromatin pulling events deforming the nucleus develop into widespread invaginations that eventually culminate into NE breakdown, we speculate that the intensity of the opposed forces on the NE increases during G2 and mitosis, and may be a driver of NE breakdown. This could happen in several ways. The centripetal pulling by chromatin could mechanically rupture the membrane and lamin nuclear surfaces through the exerted forces, or it could trigger mechanosensitive signaling cascades, as in the case of the cPLA protein (19), leading to downstream events related to different aspects of mitosis progression. Chemically, NE breakdown is known to be triggered by maturation-promoting factor (MPF), which moves into the nucleus and phosphorylates several targets (41, 42), prominently causing lamin depolymerization (25, 26, 43). The opening of the nuclear membrane is less well understood. Work in starfish indicates that it is initiated by loss of the exclusion barrier of nuclear pore complexes, followed by NE fenestration (11, 44). Recently, a mechanical action from the actin cytoskeleton has been implicated in these processes (45). Studies applying external transient tensile stress on the nuclear membrane suggest that the force range causing the typical NE deformations are sufficient to trigger nuclear membrane rupture (46).

Chromatin, through its structure and mechanics, is a key factor of nuclear function. Our results highlight that combined mechanical and/or mechano-chemical cues from condensing chromatin and cytoskeleton could also contribute to the timing and the synchronization of NE disruption with chromosome condensation during mitosis.

## Materials and Methods

### Cell culture, plasmids and transfection

HeLa cells stably expressing m-Cherry-H2B (reported previously (31)) were maintained in DMEM (Dulbecco’s Modified Eagle’s medium) with GlutaMAX (Life Technologies) supplemented with 10% (vol/vol) fetal bovine serum (FBS, Biowest), and penicillin-streptomycin (Microtech), in a humidified incubator atmosphere at 37 °C and 5% CO2.

Lipofectamine2000 (Invitrogen) was used for transfecting plasmids Emerin pEGFP-C1 (637) plasmid (Addgene ID-61993) into cells, using the protocol recommended by the manufacturer. The following day, cells were plated onto fibronectin coated glass coverslips (10 *µ*g/ml; 30 min; at 37 °C). Experiments were performed about 36-48 hours after transfection).

### Drug treatments

Calyculin A (Cell signaling technology -9902s) and latrunculin A (Sigma Aldrich-428021) were commercially purchased. Calyculin A was used at 5 nM and latrunculin A was used at 1 *µ*M concentration. Inhibitors were added to the media during the experiment, after pre-treatments acquisitions, and were maintained throughout the course of the experiment. Post calyculin A treatment, we divided the cells into two groups, named “early” and “late” phases based on their progress in rounding up and subsequent radius decrease. Cells with nuclear contour resembling that of pre-treatment are called early phase calyculin A. Cells with significantly lower nuclear radius (at least 10% less than pre-treatment) and complete deformed contour are defined late stage calyculin A. These choices are supported by time-lapse videos where the full development of the drug effect is visible (**SI Video 6**). Cells becoming rounder during the acquisition were not considered for further analysis. Latrunculin A was added to cells incubated for 16 hours with Cdk1 inhibitor (RO-3306). Videos were acquired for about 140s (see below), 20 and 50 minutes after the treatment. Mild increase in radius of G2-arrested cells compared to the regular G2 (from cell cycle analysis) is due to their prolonged arrest in G2. For Rho-associated protein kinase (ROCK) inhibition, 10 *µ*M Y27632 inhibitor was administered to cells for 30 min prior to image acquisition (**SI Video 7**). For treatments with blebbistatin, cells were treated with blebbistatin (Sigma Aldrich) at 5 *µ*M concentration for 45 min inside a dark incubator chamber to avoid photo-inactivation of the drug, then imaged for 140s. For double treatments with blebbistatin and calyculin A, cells inside a dark incubator chamber were first treated with blebbistatin (Sigma Aldrich) at 5 *µ*M concentration for 30 min and then treated with calyculin A (15 min) in presence of blebbistatin, and subsequently imaged for 140s. For cell cycle synchronisation in the G2-M transition, first cells were treated with thymidine (2 mM-Sigma Aldrich) for 14 hours, washed with PBS, released for 7 hours and then incubated further for 16 hours with Cdk1 inhibitor, RO-3306 (Seleckchem-S7747) at 10 *µ*M concentration. For ATR inhibition experiments (**SI Video 8**), 1 *µ*M of ATR inhibitor VE822 was added 2 hours prior to release from RO-3306. Cells were kept in the same inhibitor concentration throughout mitotic progression.

### Cell lysis and Immunoblotting

Cells were lysed with lysis buffer (50 mM Tris-HCl pH 8.0, 1 mM MgCl2, 200 mM NaCl, 10% Glycerol, 1% NP-40) supplemented with protease (Roche) and phosphatase inhibitors (Sigma). Cell lysates boiled with Laemmli buffer were resolved using Mini-PROTEAN® (Biorad) precast gels, transferred to 0.45 nitrocellulose membrane, and probed overnight at 4 °C with primary antibodies against pospho-Lamin A/C (Ser22) (D2B2E from CST) and vinculin (V9131 from Sigma Aldrich). After washings with 1X PBS, membranes were incubated with secondary antibodies for 1 hour at RT and acquired using ChemiDoc imaging system (Image Lab v5.0).

### Imaging and image processing

Confocal Spinning Disk microscope (Olympus) equipped with IX83 inverted microscope provided with an IXON 897 Ultra camera (Andor), Software cellSens Dimension 1.18, and attached with 100X silicone immersion objective (Refractive Index = 1.406; Numerical Aperture = 1.35) was used for HeLa cell imaging. 500 frames were acquired sequentially from Green (488 nm) and red (561 nm) channels at a maximum speed with individual exposure time of 100 ms (approx. 4 frames per second). For cell cycle based analysis, time-points were taken every 3 to 4-hour interval by acquiring 500 frames of each channel. Each cell was imaged for maximum of 5 time-points in a span of 12 hours and long-term acquisitions from the same cell was avoided to reduce the effect of phototoxicity. Cells were synchronized and released to univocally assign their cell cycle stage by monitoring their growth along the 12 hours. For treatments, multiple position acquisition was used to acquire the same cells at different time-points. Images were then processed using the ImageJ software.

Effective bending modulus and tension of the NE were obtained by fitting the fluctuation spectrum with Equation 1 for modes 6-34. Modes below 5 were excluded because influenced by the cell shape (23) and higher modes above 34 were affected by noise due to the acquisition exposure time. From fluctuation dynamics, relaxation time of mode 3 was obtained by fitting the autocorrelation function of the fluctuation amplitudes for mode 3 with a single exponential (3). Invaginations were first identified by change in contour fluctuations and confirmed by looking at videos. The mean of the contour shape for the first 10 frames was subtracted from the contour of each frame as reference. The depth is the difference between the steady state contour and the minimum of the invagination. The width is determined by the points corresponding to 10% of the depth.

### Convention for Fourier transform in the flickering code

Equation (1) is derived in (3) and uses the following non-unitary convention for the 2D Fourier transform of the displacement function 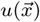:

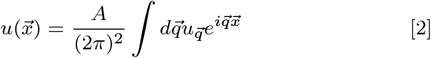

and the inverse transform is

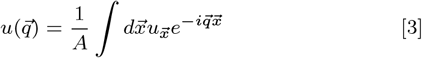

where *A* = *L* × *L* is the area of the membrane.

In order to match the Fourier transform with the discrete Fourier series calculated in Matlab (Fast Fourier Transform, FFT), the Fourier coefficients coming out from Matlab’s FFT of *u* need to be corrected by:

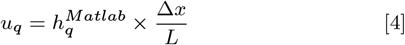

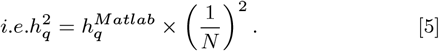

## ACKNOWLEDGMENTS

The authors thank Paolo Maiuri, Giorgio Scita, Nishit Srivastava and Matthieu Piel for useful feedback on this manuscript, and Nils Gauthier for sharing reagents. M.C.L. is supported by the Italian Association for Cancer Research (AIRC), grant AIRC-IG (REF: 23258). P.C. is supported by the Engineering and Physical Sciences Research Council (EPSRC) (EP/R011443/1). Work in M.F.’s laboratory was supported by grants from AIRC, Telethon-Italy, Ministero dell’Istruzione, dell’ Università e della Ricerca and the European Commission. V.I. was funded by the EPSRC and Sackler scholarships, and by the Wellcome Trust Junior Interdisciplinary fellowship. G.R.K. was supported by Marie Curie Initial Training Networks (ITN), (FP7 ‘aDDRess’) fellowship and Italian Association for Cancer Research (AIRC) fellowship.

