## Supplementary Information for "Centripetal nuclear shape fluctuations associate with chromatin condensation towards mitosis"

**Tables** **2-4**

**Figures** **5-14**

**Videos** **15**

**Table S1**. Fluctuation properties of the nuclear envelope (NE) across the cell cycle

| Cell cycle | RBCs | G1 | S | G2 | ME | ML |
|---|---|---|---|---|---|---|
| Number of cells | 31 | 20 | 15 | 19 | 9 | 11 |
| Effective bending modulus ($10^{-20}$ J) | 12.5 ± 0.8 | 2.3 ± 0.7 | 1.4 ± 0.4 | 2.2 ± 1.2 | 2.2 ± 0.5 | 1.3 ± 0.4 |
| Effective tension ($10^{-7}$ N/m) | 12.0 ± 0.8 | 17.1 ± 2.2 | 23.6 ± 2.4 | 19.5 ± 2.7 | 3.9 ± 0.4 | 1.8 ± 0.5 |
| Radius (µm) | 4.18 ± 0.04 | 7.5 ± 0.1 | 8.5 ± 0.3 | 11.1 ± 0.5 | 9.6 ± 0.3 | 9.2 ± 0.4 |
| Relaxation time mode 3 (s) | 0.035 ± 0.008 | 2.1 ± 0.1 | 2.4 ± 0.1 | 2.0 ± 0.2 (14 cells) | 2.4 ± 0.2 | 2.9 ± 0.2 |

Nuclear radius, effective tension, effective bending modulus and fluctuation timescale of mode 3, measured by flickering spectrometry, were compared at different times during the cell cycle. 6-34 modes were considered for the HeLa cell cycle, modes 8-20 for RBCs. Measurements were consolidated from 3 independent experiments for each phase. The nuclear radius is the mean calculated from the center of the nucleus. The average values and the respective standard errors of the mean (SEM) were calculated from the number of cells indicated in the table. Pairwise statistical comparisons were performed using the two-sided Mann-Whitney U test. Data shown in **Fig. 1d-g**.

**Table S2.** P values < 0.05 for Figures in the manuscript

| **Fig. 1** | | **Fig. 2** | | **Fig. 3** | |
|---|---|---|---|---|---|
| **Effective tension ($10^{-7}$ N/m)** | **P values** | **Effective tension ($10^{-7}$ N/m)** | **P values** | **Skewness** | **P values** |
| G1 – ME | 2.45 x $10^{-5}$ | Calyculin A early | 0.0142 | G1-G2 | 0.0282 |
| G1 – ML | 1.95 x $10^{-5}$ | Calyculin A late | 1.65 x $10^{-4}$ | G1-ME | 1.03 x $10^{-4}$ |
| S – G2 | 0.01 | Latrunculin A 20 min post-treatment | 8.7 x $10^{-5}$ | G1-ML | 6.04 x $10^{-6}$ |
| S – ME | 3.47 x $10^{-4}$ | Latrunculin A 50 min post-treatment | 3.0 x $10^{-4}$ | S-ME | 0.0079 |
| S – ML | 4.75 x $10^{-8}$ | **Radius (μm)** | **P values** | S-ML | 6.25 x $10^{-5}$ |
| G2 – ME | 4.79 x $10^{-5}$ | Calyculin A late | 0.0024 | G2-ML | 2.81 x $10^{-4}$ |
| G2 – ML | 5.96 x $10^{-6}$ | | | ME-ML | 0.0079 |
| ME – ML | 0.005 | | | Calyculin A Control -ME | 0.0021 |
| **Radius (μm)** | **P values** | | | Calyculin A Control -ML | 2.31x $10^{-5}$ |
| G1 – S | 0.0044 | | | VE822 | 0.0265 |
| G1 – G2 | 1.4 x $10^{-7}$ | | | | |
| G1 – ME | 2.45 x $10^{-5}$ | | | | |
| G1 – ML | 2.58 x $10^{-4}$ | | | | |
| S – G2 | 7.53 x $10^{-5}$ | | | | |
| S – ME | 0.015 | | | | |
| G2 – ME | 0.04 | | | | |
| G2 – ML | 0.02 | | | | |
| **Relaxation time mode 3 (s)** | **P values** | | | | |
| G1 – ML | 0.0027 | | | | |
| S – ML | 0.0293 | | | | |
| G2 – ML | 0.0031 | | | | |

P values reported for data shown in **Fig. 1-3**. Pairwise statistical comparisons were performed using the two-sided Mann-Whitney U test.

**Table S3.** Biophysical properties of NE upon biochemical perturbations

| Treatment | Number of cells | Tension ($10^{-7}$ N/m) | Radius (µm) | Relaxation time mode 3 (s) |
|---|---|---|---|---|
| Calyculin A pre-treatment | 21 | 13.3 ± 1.7 | 8.2 ± 0.1 | 2.1 ± 0.2 (5 cells) |
| Calyculin A early | 9 | 7.2 ± 1.1 | 7.9 ± 0.2 | 2.7 ± 0.1 (5 cells) |
| Calyculin A late | 12 | 2.6 ± 0.4 | 6.3 ± 0.3 | |
| Latrunculin A pre-treatment | 13 | 9.9 ± 1.4 | 12.1 ± 0.4 | |
| Latrunculin A 20 min | 11 | 2.9 ± 0.4 | 11.4 ± 0.4 | |
| Latrunculin A 50 min | 9 | 2.2 ± 0.3 | 11.6 ± 0.5 | |

Nuclear radius and tension measured by flickering spectrometry upon calyculin A and latrunculin A perturbations. Two biological replicates were measured for each condition, and the same nuclei were recorded before and after biochemical treatments. The relaxation time of mode 3 is reported in the case of calyculin A treatment. The average values and the respective standard errors of the mean (SEM) were calculated from the number of cells indicated in the table. Data shown in **Fig. 3**.

**Figure S1.** Nuclear (cross-section) area increases throughout the cell cycle in Hela cells

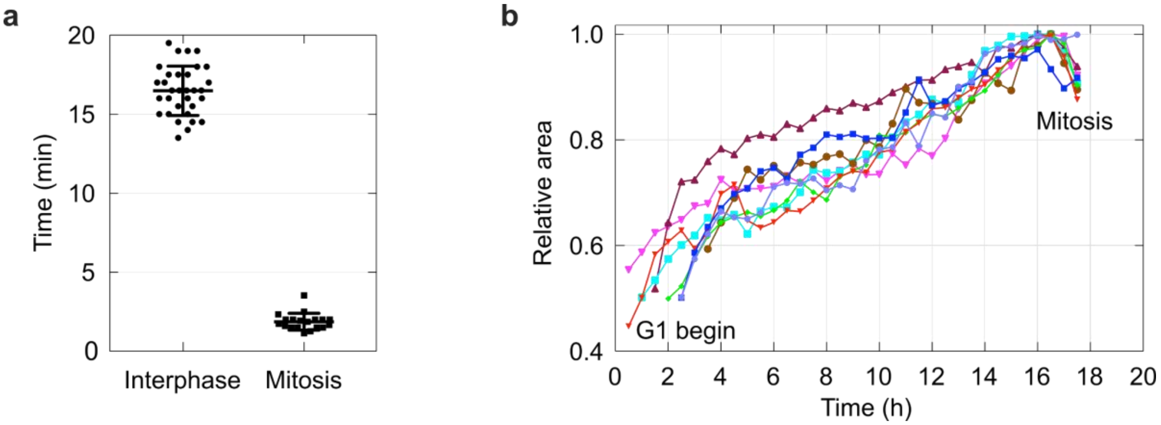


Hela cell nuclei are tracked using the H2B-mCherry marker. (**a**) Average time taken by cells to complete one cell cycle, separately in interphase (after formation of daughter nuclei post mitosis) and in mitosis (from beginning of condensation to formation of daughter nuclei). Interphase: -16.48 ± 0.2754, n=32; mitosis: -1.75 ± 0.1118, n=10. (**b**) Relative growth of nuclear cross section area across cell cycle. Observed a continuous increase in nuclear area throughout the cell cycle reaching maximum at late G2 and with a minor reduction at the onset of mitosis (n=8). This is in agreement with data on nuclear radii in **Fig. 1d**.

**Figure S2.** Nuclear shape properties of the same cells throughout their cycle

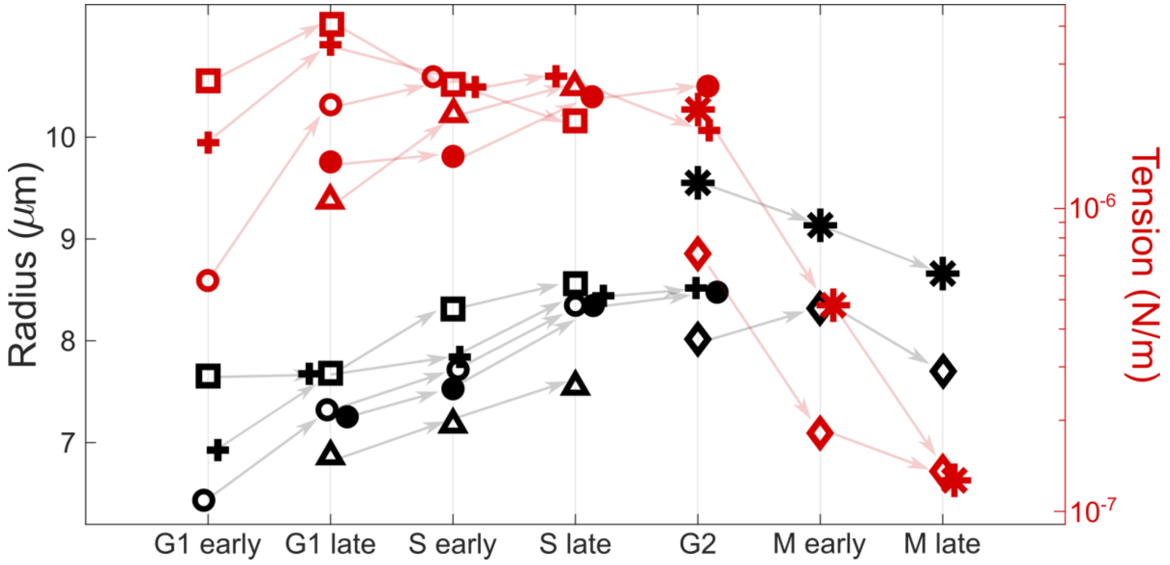


Seven cells followed during their development towards mitosis starting either from arrest in G1 or late G2. Their behaviour is generally in agreement with the bulk shown in **Fig. 1**.

**Figure S3.** Autocorrelation plot of the temporal evolution of mode 3 from the static spectrum

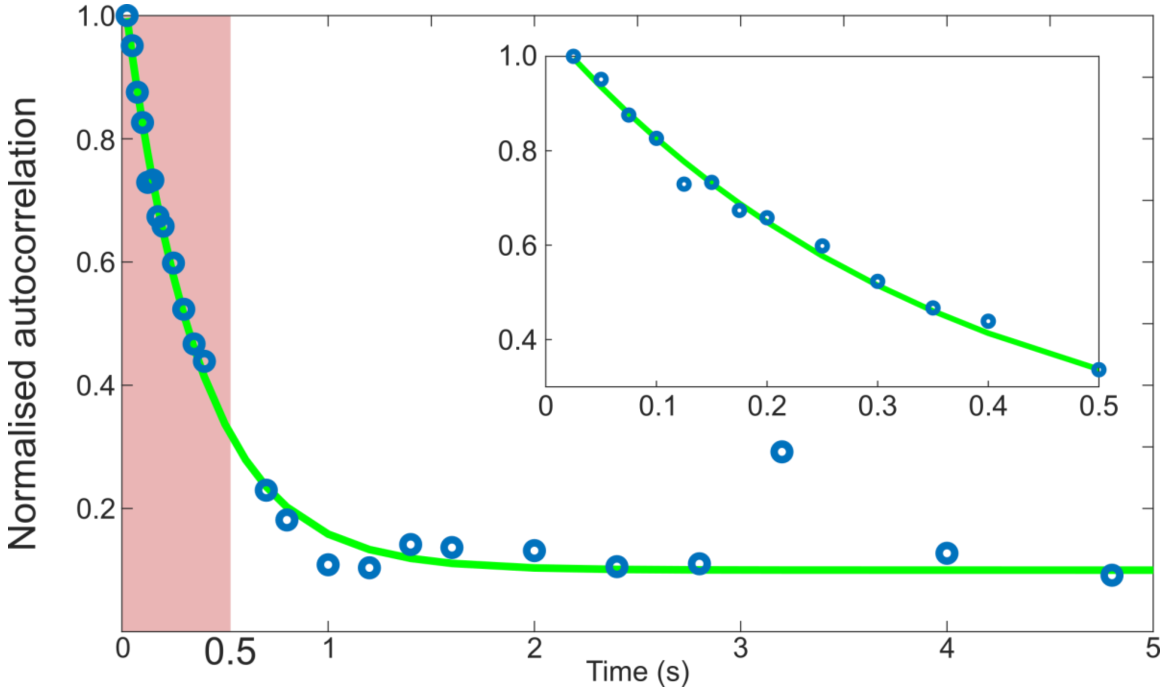


Typical autocorrelation function of the fluctuation amplitudes for mode 3 as function of time. This function has been fitted for times below 0.5 s with a single exponential to obtain the relaxation time of nuclei, as suggested by theory (Yoon *et al*. 2009 [ref 3 main text]).

**Figure S4.** Calyculin A treatment does not affect Lamin A/C phosphorylation

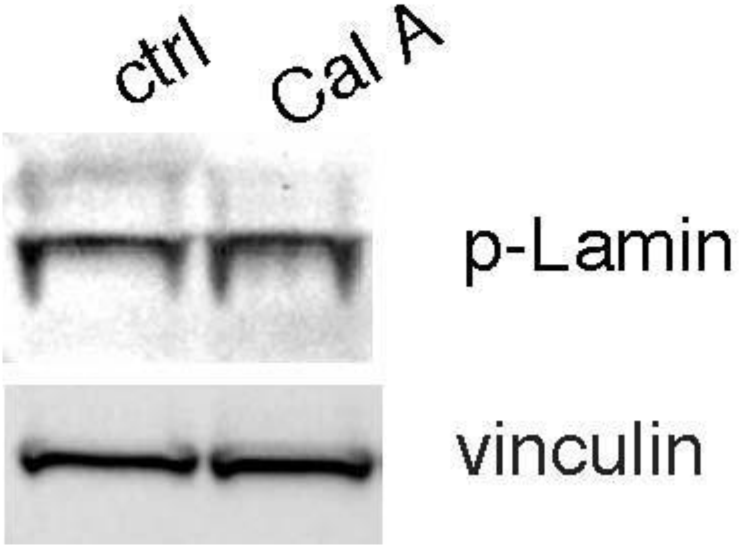


Western blot analysis of control HeLa cells and HeLa cells treated with 5 nM calyculin A for 20 minutes, probed for phospho-Lamin A/C (Ser22) and Vinculin. Calyculin A treatment does not result in changes in the level of Lamin A/C phosphorylation.

**Figure S5.** Control experiment: actomyosin contractility does not cause nuclear invaginations

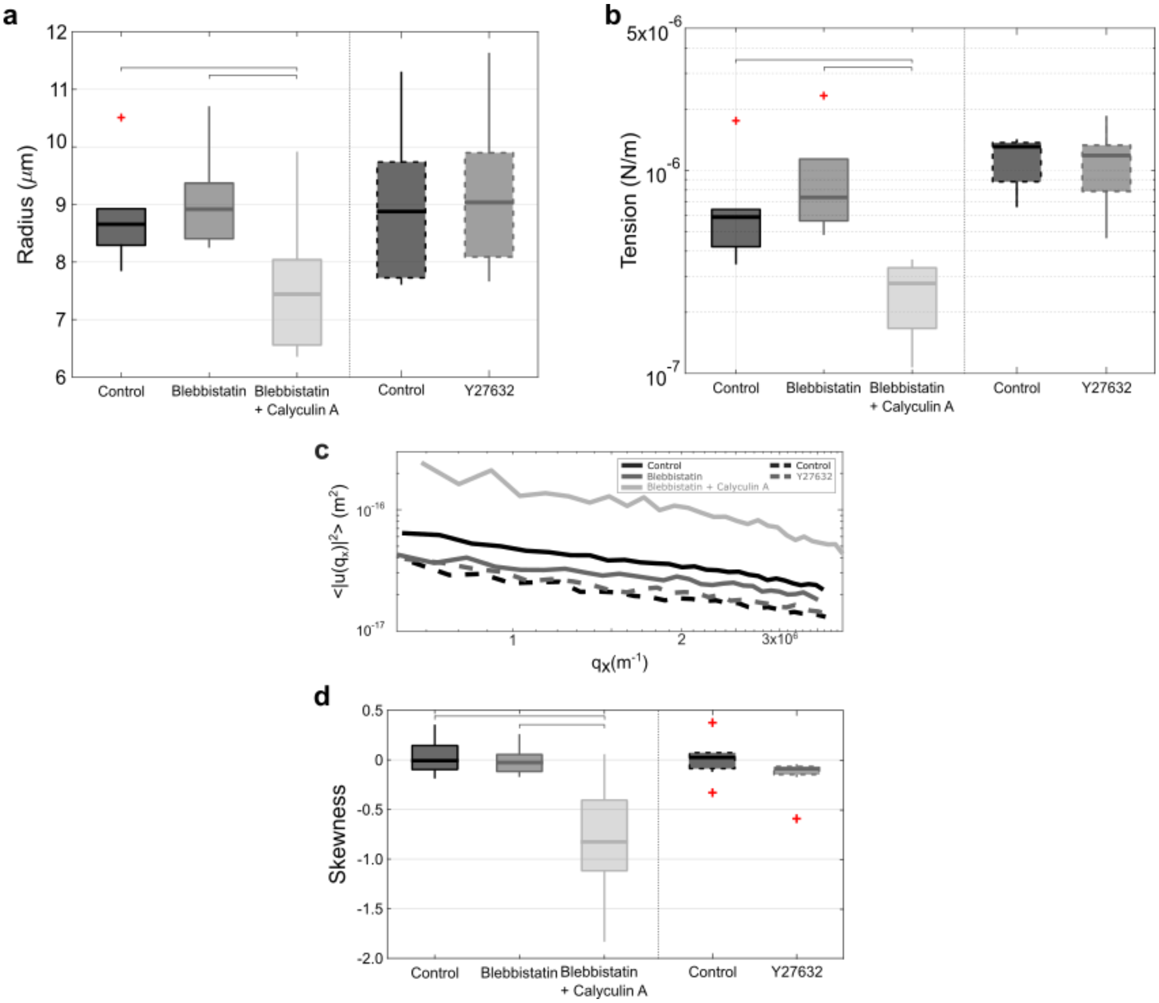


Experiments to rule out the role of myosin-2 mediated contractility as origin of nuclear invaginations due to calyculin A treatment. Blebbistatin and Y27632 (contrary to blebbistatin which is inactivated by blue light, Y27632 is not affected by illumination) confirm that the invagination phenotype is independent of actomyosin contractility. Nuclear radius, tension, amplitudes of nuclear envelope fluctuations and skewness of fluctuation distribution measured for 8 unsynchronised cells before and after blebbistatin and Y27632 treatments of 30 min (in two separate experiments) and blebbistatin + calyculin A 45min treatment (first 30 min incubation with only blebbistatin + 15 min incubation with both). No significant difference for all the properties measured between control and blebbistatin/Y27632 treatments, while the difference is significant with respect to the combination of blebbistatin + calyculin A as expected from **Fig.3**. P values highlighted in the figure: radius 0.0493 (control - blebbistatin + calyculin A) and 0.0295 (blebbistatin - blebbistatin + calyculin A); tension 0.0027 (control - blebbistatin + calyculin A) and 0.0016 (blebbistatin - blebbistatin + calyculin A); skewness 0.0174 (control - blebbistatin + calyculin A) and 0.0035 (blebbistatin - blebbistatin + calyculin A).

**Figure S6.** Shape of Invaginations in early mitosis is compatible with prediction of a pinning force

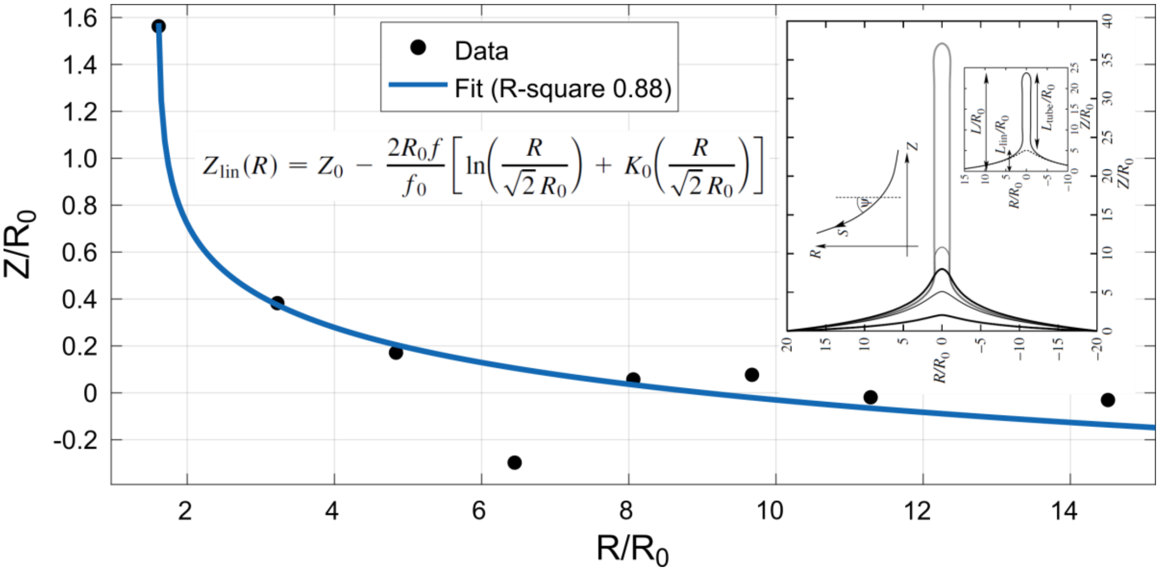


The shape of invaginations at the early stage of nuclear mitosis resemble the shape of an emerging tube when the membrane is pulled by a point force f. The data for one side of the invagination are fitted with the equation in the inset formulated by Derényi *et al*. 2002 [ref 30 main text], by knowing $f_0$ and $R_0$, which are related to the bending modulus and tension of the membrane (in our case effective bending modulus and tension).

**Figure S7.** Absence of separation between NE and chromatin globule surface (CGS) at the invagination sites

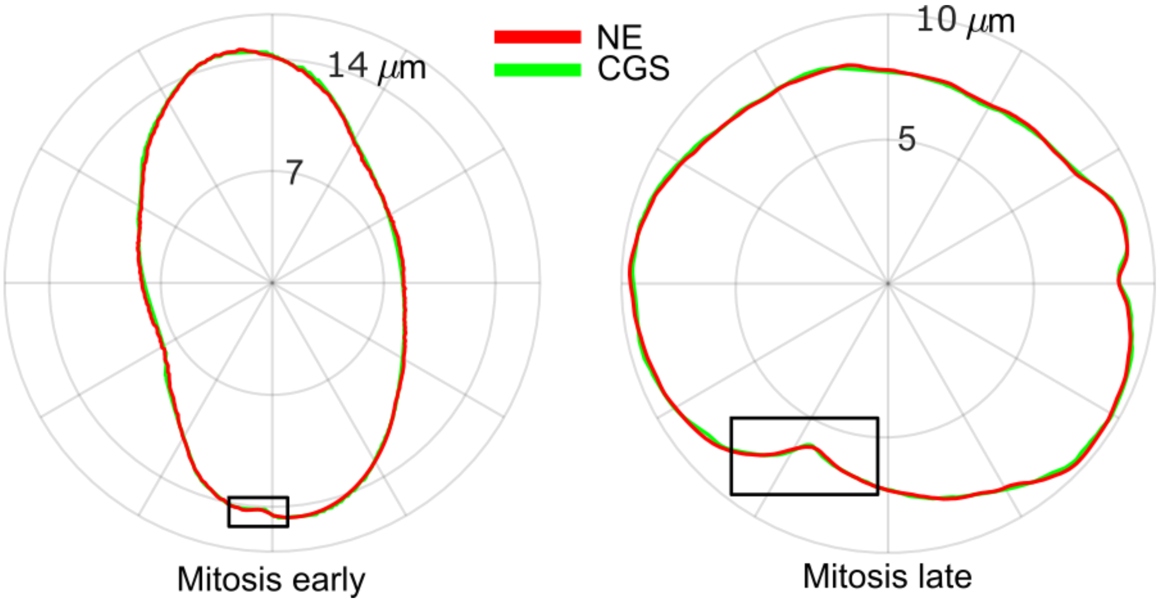


Instantaneous contours of NE and CGS for representative cells in early and late mitosis do not exhibit any separation in the site of invaginations (highlighted in the black square). No separation between NE and CGS was reported throughout the invagination period for all cells analysed.

**Figure S8. Correlation between histones and transient invaginations during mitosis**

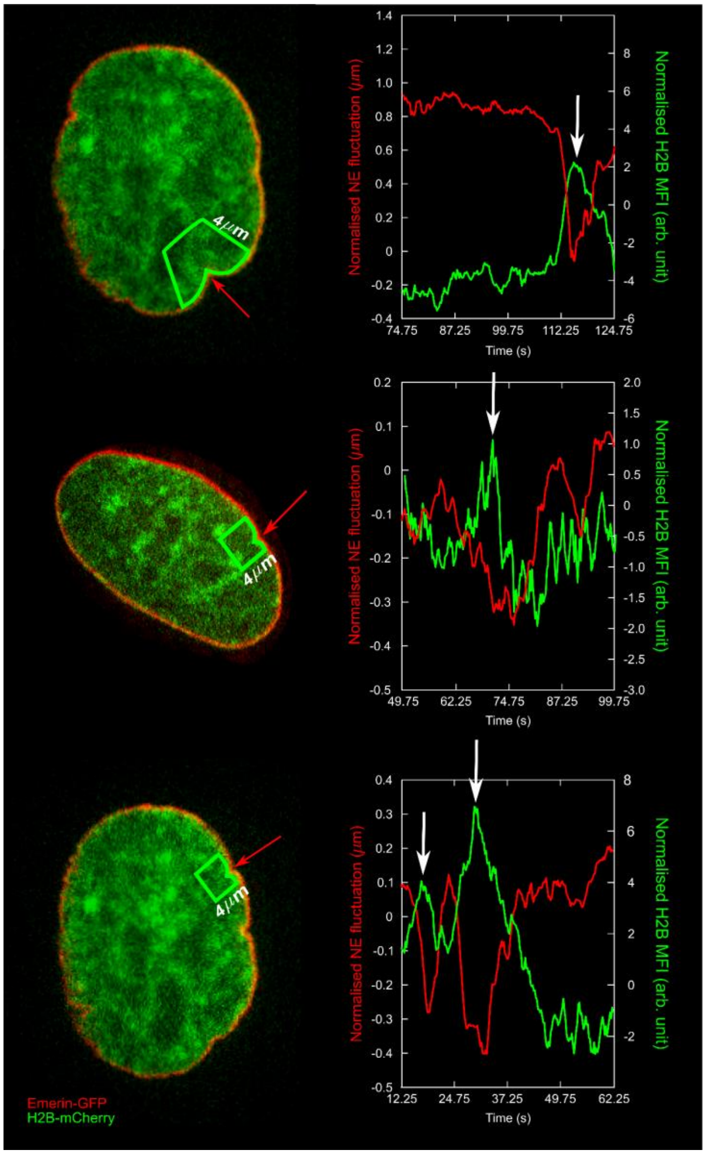


Examples of the negative correlation between the fluorescence signals of histones (green) in the proximity of the invagination (area highlighted in the green in the image of the nuclei) and the relative NE deformation (red). The green plots show the mean fluorescence intensity of the area of interest normalised for the mean fluorescence intensity of the entire nuclei during transient invaginations, while the red plots show the nuclear contour for the angle at which the invagination is at its maximum and subtracting the initial frame. As shown in **SI Video 9**, the third nucleus has repetitive invaginations in the same section of the contour, well described by the relative NE deformations and chromatin fluorescence trends.

**Figure S9.** Effect of ATR inhibition on NE

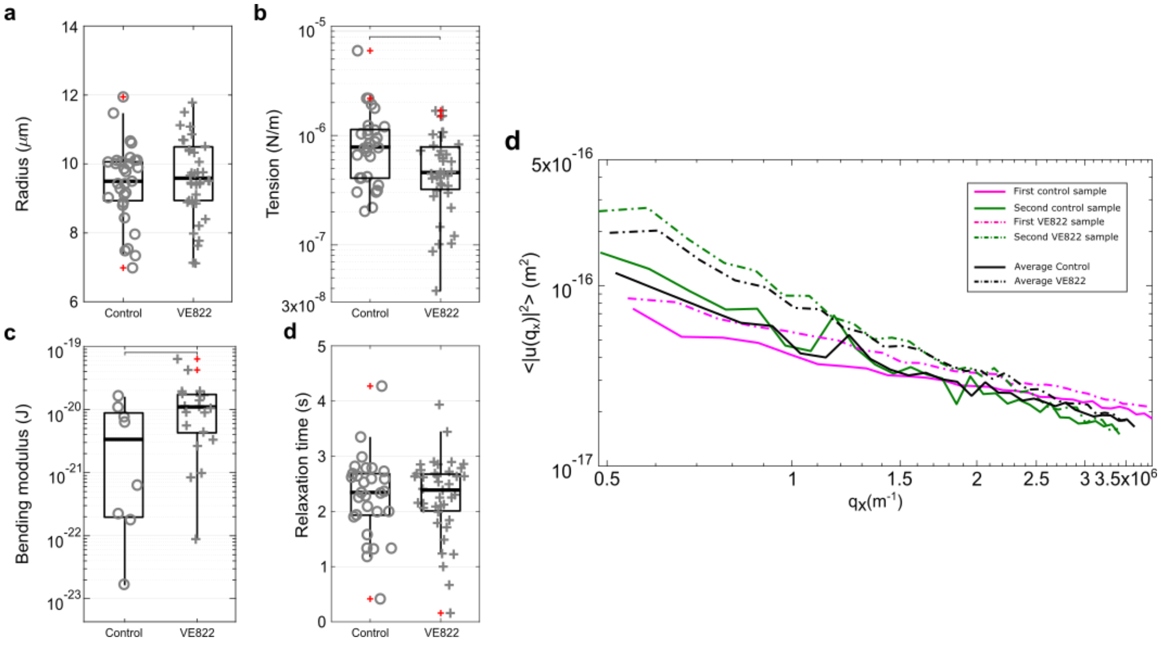


Effective tension is reduced (p value = 0.0218) and effective bending modulus increases (p value = 0.0338) in cells treated with VE822. The increase of bending modulus could be due to changes in the lipid composition as suggested by Kidiyoor *et al.* 2020 [ref 32 main text]. The radius and the relaxation time for mode 3 do not show significant difference before and after ATR inhibition. Data from 2 sets of experiments: 29 cells for control, 38 cells for VE822 treatment.

**Figure S10.** Summary of nuclear contour fluctuations throughout the cell cycle and after chemical perturbations

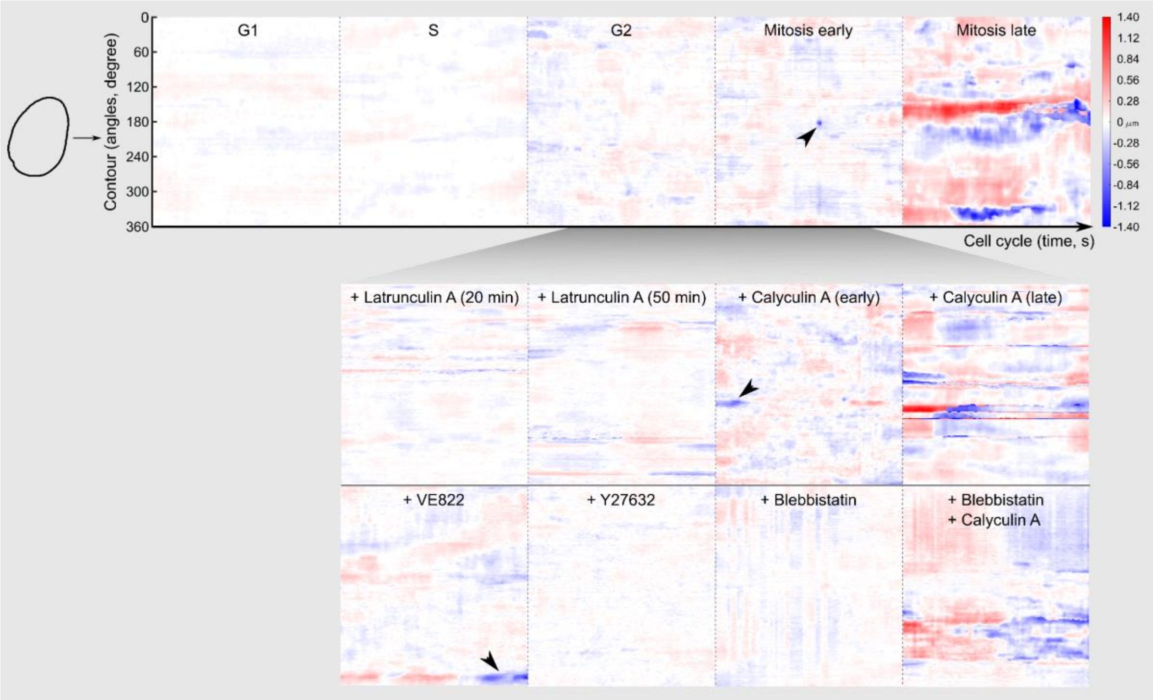


Heat maps highlighting the contour fluctuations of representative nuclei at different times of the cell cycle and after treatments. For each nucleus, the x axis indicates the duration of the recorded video (500 frames = 125 s), the y axis the contour profile angle, and the color map is the deviation from the mean contour (negative inward, positive outward). Black arrows indicate transient and localised invaginations that characterise early mitosis.

**SI Video 1.** Nuclear shape fluctuations in interphase (G1,S,G2) nuclei.

**SI Video 2.** Nuclear shape fluctuations in mitotic (early and late) nuclei. NE labeled with Emerin-GFP and chromatin labeled with H2BmCherry. Scale bar =5μm

**SI Video 3.** Nuclear shape fluctuations in calyculin A treated (early and late) nuclei.

**SI Video 4.** Nuclear shape fluctuations in latrunculin A treated (early and late) nuclei.

**SI Video 5.** Nuclear shape fluctuations in blebbistatin and calyculin A treated nuclei.

**SI Video 6.** Entire mitosis process from early mitosis to nuclear envelope breakdown (total recording of 20 minutes at 5 seconds/frame rate).

**SI Video 7.** Nuclear shape fluctuations of ROCK inhibited-nuclei treated with Y27632.

**SI Video 8.** Correlation between the fluorescence signal of histones and NE deformation during transient invaginations

**SI Video 9.** Nuclear shape fluctuations of ATR inhibited-nuclei arrested in mitosis and treated with VE822.

**SI Video 10.** Representative 3D kymograph of nuclear shape fluctuations during mitosis.
